# Inhibition of PIKfyve kinase induces senescent cell death by suppressing lysosomal exocytosis and leads to improved outcomes in a mouse model of idiopathic pulmonary fibrosis

**DOI:** 10.1101/2025.03.19.644224

**Authors:** Anna Barkovskaya, Kristie Kim, Apoorva Shankar, Gabriel Meca-Laguna, Michael Rae, Claude Jourdan Le Saux, Amit Sharma

## Abstract

Cellular senescence is a pivotal hallmark of aging, which limits lifespan and contributes to the development of age-related diseases. Efforts to identify senolytics - drugs that selectively eliminate senescent cells, have so far yielded candidates with limited translational potential. Here, we characterize the senescent cell surface proteomic landscape and identify proteins that are abnormally present on the plasma membrane of senescent cells. Many of these proteins are lysosomal enzymes, pointing to lysosomal exocytosis as a likely mechanism that leads to their persistent display on the cell surface. Blocking lysosomal exocytosis via PIKfyve kinase inhibition with a small molecule drug apilimod results in selective killing of senescent cells *in vitro*, while this treatment does not affect quiescent and proliferating cells. Furthermore, apilimod can be safely administered *in vivo* and effectively removes senescent cells and reduces tissue remodeling in a bleomycin mouse model of pulmonary fibrosis. We conclude that apilimod is an effective and well-tolerated senolytic that may be useful for the treatment of senescence-associated diseases of aging.

## Introduction

Senescent cells act as a unifying link among the hallmarks of aging, serving both as consequences and as drivers of age-related damage (1, 2). These cells arise from genomic instability, telomere attrition, and epigenetic alterations, which they further exacerbate through persistent DNA damage signaling and the senescence-associated secretory phenotype (SASP) (3). They contribute to mitochondrial dysfunction by generating reactive oxygen species (ROS), impairing energy metabolism, and disrupting proteostasis through inefficient protein clearance and SASP-mediated extracellular matrix degradation (4). Additionally, senescent cells disturb nutrient-sensing pathways, such as mTOR and insulin/IGF-1 signaling, and jeopardize stem cell niches by releasing inflammatory factors that induce stem cell exhaustion (5). Their SASP modifies intercellular communication, drives chronic inflammation, also referred to as “inflammaging,” and further propagates aging-related dysfunction. Importantly, senescent cells create self-reinforcing feedback loops that amplify other hallmarks, establishing them as central drivers of systemic aging (6). Therapeutic strategies that target senescent cells, such as senolytics and senomorphics, have shown promise in alleviating age-related pathologies, emphasizing their pivotal role as a common link in aging (7).

Despite advancements in therapies targeting senescent cells, there is an urgent need to discover new and improved strategies that can overcome the complexities of senescence biology and the limitations of current methods. For senolytic interventions, immense potential remains untapped. Furthermore, the heterogeneity of senescent cells across tissues and disease contexts (8, 9) and the physiological roles of these cells (10) highlight the necessity for a more varied arsenal of therapies tailored to specific senescent cell populations, SASP profiles, and biological contexts.

The lack of strong surface markers for senescence presents a significant obstacle to investigating the biology of senescent cells and developing effective therapies to eliminate them. In addition to enabling the differentiation of non-senescent cells from their senescent counterparts *in vitro* and in aged and diseased tissue samples, identifying and validating previously unrecognized markers on senescent cell surfaces could reveal promising targets for the selective removal of senescent cells by engineered immune cells and improve senolytic drug targeting.

Here, we leveraged the innovative Ligand-Receptor Capture (LRC) TriCEPS technology (11) to identify senescent cell surface proteins in an unbiased manner. This led to the discovery of several novel senescent cell surface markers, several of which are lysosomal proteins — most notably the transmembrane protein lysosomal-associated membrane protein 1 (LAMP-1) (12). Alongside this, we found evidence of plasma membrane damage and release of lysosomal enzymes into the medium, consistent with senescent cells being in a chronic state of lysosomal exocytosis. We showed that senescent cells’ reliance on this process can be therapeutically exploited by pharmacological inhibition. Inhibitors of phosphoinositide kinase, FYVE-type zinc finger containing (PYKfyve kinase) selectively induce cell death in senescent cells through a nonapoptotic form of cell death that appears to be methuosis, a distinctive form of cell death characterized by high levels of vacuolization.

To validate our findings in a model of age-related disease, we induced interstitial lung remodeling in mice to mimic idiopathic pulmonary fibrosis (IPF). IPF is a complex and debilitating lung disease characterized by progressive and irreversible scarring (fibrosis) of the lung parenchyma (13). Surveillance and reporting methods vary internationally, but it is evident that the incidence of IPF rises with age, and the mean age of an IPF patient is approximately 65 years old (14, 15). Numerous studies have reported an increased burden of senescent cells (epithelial, mesenchymal, and endothelial) in the lungs of IPF patients compared to healthy individuals. Animal models of pulmonary fibrosis, especially mice, have provided evidence supporting the causal role of cellular senescence in fibrotic lung diseases (16, 17). In this study we used a mouse model of IPF to confirm the senolytic activity of apilimod dimesylate, positioning it as a potential therapeutic intervention.

## Materials and Methods

### Cell culture

IMR-90 fibroblasts (ATCC, USA: CCL-186) were maintained at 37°C in humidified air with 5% CO_2_ and 3% O_2_ in Dulbecco’s Modified Eagle’s Medium (DMEM) (Corning; 10-013-CV) supplemented with 10% Fetal Bovine Serum (FBS) (Millipore Sigma, USA; F4135) and 1X Penicillin–Streptomycin (Corning; 30-001-CI). Cumulative population doubling (PD) was calculated using the following equation: PD= P-PD + (log (H / S)/ log 2), where H is the number of cells at harvest, and S is the number of cells seeded; P-PD is the PD number in the previous cell collection. Primary human endothelial cells (EC) were purchased from the Coriell Institute for Medical Research (AG10770). EC were maintained in promo cell basal medium MV2 (PromoCell; C-22221) supplemented with Growth Medium MV 2 Supplement Pack (PromoCell; C-39221) and used for up to 10 passages. Quiescence was induced by replacing culture media with media containing 0.2% FBS. All cells were routinely tested for mycoplasma. The following reagents were used for treatment: apilimod (MedChem Express, HY-14644); apilimod dimesylate (ApiD) (Axon Med Chem AXON 2500); vacuolin-1 (EMD Millipore 67300); Z-VAD-FMK (Selleck Chemicals S7023).

### Senescence induction

To generate doxorubicin-induced senescent (S-dox) cells, IMR-90 cells were treated with 300 nM of doxorubicin hydrochloride (Millipore Sigma, USA; Cat# 504042) for 24 hours. After that, the media was replaced to remove doxorubicin. For radiation-induced senescent (S-IR) cells, the cells were irradiated with 10 Gy X-ray and maintained in culture. For S-dox endothelial cells (S-dox EC), primary human EC were treated with 250 nM of doxorubicin in PromoCell basal medium MV2, supplemented with Growth Medium MV2 Supplement Pack, for 24 hours. Senescent cells were assayed on day 10 following treatment. Replicative senescence (RS) was achieved by continuous culture until proliferation was exhausted at PD ≥ 58. RS was confirmed using the SA-β-gal staining method described below. For non-senescent (NS) controls, proliferating cells with PD ≤ 35 were used.

### Senescence-associated ß-galactosidase staining

Senescence-associated ß-galactosidase (SA-β-gal) activity assay was performed as previously described (18), using the Senescence Detection Kit (BioVision; K320) and following the manufacturer’s instructions. During the staining process, cells were incubated for 24 hours at 37°C without CO_2_ and subsequently imaged using bright-field microscopy.

### qPCR gene expression

Total RNA was isolated from cell pellets using the Quick-RNA MiniPrep (Zymo Research; R1055), following the manufacturer’s instructions. cDNA synthesis was performed using the PrimeScript RT Master Mix (Takara; RR036B), in accordance with the manufacturer’s guidelines. Quantitative PCR was conducted on a StepOnePlus™ Real-Time PCR System (ThermoFisher) using primers and probes obtained from Applied Biosystems TaqMan Gene Expression assays and TaqMan Fast Advanced Master Mix (Applied Biosystems; 4444557). Relative mRNA expression was evaluated using the comparative threshold (Ct) method, normalizing target cDNA Ct values to those of actin. All reactions were carried out in triplicate. The following primer-probes were used: LMNB – HS01059210_m1; CDKN2A – HS00923894_m1; CDKN1A – HS00355782_m1; IL6 – HS00174131_m1; IL8 – HS00174103_m1; IL1α – HS00174092_m1.

### Proliferation assay

Cell proliferation was measured by incorporating EdU in dividing cells using the Click-iT™ EdU Cell Proliferation Kit for Imaging, Alexa Fluor 488 dye (Thermo Scientific; C10337), following the manufacturer’s instructions. Briefly, cells were cultured with EdU (10 μM) for 24 hours, fixed in 4% PFA for 10 minutes, washed in PBS, and permeabilized with 0.5% Triton X-100 for 15 minutes. Cells were then washed and incubated in Click-iT® reaction cocktail, and DNA was stained with Hoechst 33342 (Invitrogen; Cat# H3570). For senescent cells, fibroblasts were plated one day before induction of senescence. Staining was performed 7 days after senescence induction. EdU incorporation was visualized and imaged using a fluorescence microscope. The percentage of EdU-positive cells was quantified by counting the total number of cells and EdU-positive cells over 12 microscopic fields per assay.

### Immunofluorescence

Cells were seeded in black-wall, glass-bottom 96-well plates (Ibidi; 89626). Following treatments, they were fixed with 4% PFA (Thermo Scientific; AAJ19943K2) in PBS for 15 minutes at room temperature. For intracellular labeling, cells were permeabilized with 0.5% Triton X-100 for 10 minutes. This step was excluded for labeling cell membrane markers. Samples were blocked with 5% goat serum (GS) (Gibco, 16210-064) for at least 1 hour and stained with primary antibodies overnight at 4°C on a shaker. The following primary antibodies were used: anti-γ-H2AX (pSer139) (Novus Biologicals; NB100-74435), anti-HMGB1 (abcam; ab18256), anti-cathepsin C (CATC) (Santa Cruz; sc-74590), anti-TPP1 (Millipore; MABW1806), anti-CA2D1 (CACNA2D) (Thermo Fisher; MA3-921), anti-LAMP1 (Santa Cruz, clone H4A3). Samples were washed with PBS and stained with secondary antibodies (1:1000 Alexa Fluor 488 goat anti-mouse (Invitrogen; A11029); Alexa Fluor 546 goat anti-rabbit (Invitrogen; A11010)) in 5% GS-PBS for 1 hour, then washed and stained with Hoechst 33342 for 15 minutes at room temperature. Images were taken using the EVOS M5000 microscope (Invitrogen). The fluorescence intensity was quantified using ImageJ software, measuring intensity in the area occupied by the cell and subtracting background intensity in the same image. At least five cells per image were quantified. All experiments were conducted with at least three biological replicates.

### Surfaceome-TriCEPS^TM^ ligand-receptor capture (LRC-TriCEPS)

We used the Dualsystems Biotech AG TriCEPS^TM^ V.3.0 kit to capture proteins on the surface of living cells (Cat. Number: P0529) and to identify surface proteins enriched in S-dox and S-IR IMR-90 cells according to the manufacturer’s instructions. Samples of S-dox, S-IR, and NS cells, each containing 24×10^6^ cells, were scraped into PBS (pH 6.5) to detach the cells, then washed and cooled to 4°C. Biological triplicates of all samples were prepared, and subsequent steps were performed at 4°C. Sodium metaperiodate (1.5 mM) was added to mildly oxidize the cell surface proteins, and the samples were incubated at 4°C in the dark for 15 minutes with gentle rotation. The cells were washed twice at 300xg for 5 minutes, then resuspended in 20 ml of surfaceome buffer. For surfaceome labeling, quenched TriCEPS V. 3.0 was added to the oxidized cells and incubated in the dark for 90 minutes on a rotator. The cells were then centrifuged at 4°C and stored at −80°C before further analysis. Proteomics was conducted at Dualsystems Biotech AG, where samples underwent purification using solid-phase chromatography, stringent washing to remove nonspecific interactions, reduction, alkylation, and digestion with trypsin. The tryptic peptides were collected for LC-MS/MS analysis. Mass spectrometry was performed on a Thermo Orbitrap Elite spectrometer equipped with an electrospray ion source. Tryptic peptides were analyzed in data-dependent acquisition mode (TOP20) over an 80-minute gradient, utilizing a 15 cm C18 packed column.

### Surfaceome-TriCEPS^TM^ data analysis

Progenesis software was used to align raw files and to detect features. The Comet search engine was employed for spectrum identification. The Trans-Proteomic Pipeline was used to statistically validate putative identifications and protein inference. Upon inferring proteins, relative quantification of samples was performed based on ion-extraction intensity, and differential protein abundance was assessed using a statistical ANOVA model followed by multiple testing corrections. This model assumes that measurement error follows a Gaussian distribution, considers individual features as replicates of a protein’s abundance, and explicitly accounts for this redundancy. It evaluates each protein for differential abundance across all pairwise comparisons and reports the p-values. Subsequently, p-values are adjusted for multiple comparisons to control the experiment-wide false discovery rate (FDR). The adjusted p-value (q-value) for each protein is plotted against the magnitude of the fold enrichment between the two experimental conditions.

The quantitative differences between perturbations are shown as a volcano plot. The x-axis represents the mean ratio fold change (on a log2 scale). The y-axis represents the statistical significance p-value of the ratio fold change for each protein (on a −log10 scale) (adjusted p value).

Pathway enrichment analysis was performed using Metascape. To identify the differentially regulated pathways, all statistically enriched terms were identified, and accumulative hypergeometric p-values and enrichment factors were calculated and used for filtering. The remaining significant terms were hierarchically clustered into a tree based on Kappa-statistical similarities among their gene memberships. Then, a 0.3 kappa score was applied as the threshold to cast the tree into term clusters.

### Immunoblotting and isolation of cell membrane and cell surface proteins

Cells were lysed in RIPA buffer (20 mM Tris-HCl pH 7.5, 150 mM NaCl, 1 mM Na_2_EDTA, 1 mM EGTA, 1% NP-40, 1% sodium deoxycholate, 2.5 mM sodium pyrophosphate, 1 mM beta-glycerophosphate, 1 mM Na_3_VO_4_, and 1 µg/ml leupeptin) containing a protease inhibitor cocktail (Cell Signaling Technology; 5871). Cell suspensions were incubated on ice for 10 minutes, followed by microcentrifugation at 4°C for 15 minutes to clarify the lysate of cell debris. To isolate cell membrane and surface proteins, cell lysates were prepared using the Mem-PER Plus Membrane Protein Extraction Kit (Thermo Scientific; 89842) or the Pierce Cell Surface Protein Isolation Kit (Thermo Scientific; 89881). Protein concentration was quantified using the BCA Protein Assay (Thermo Scientific; 23227). Equal amounts of protein (10-20 µg) were separated by SDS/PAGE and transferred onto PVDF membranes (Bio-Rad; 1620177) using semi-dry transfer. Membranes were incubated with primary antibodies overnight at 4°C, followed by an 1 hour incubation with secondary antibodies at room temperature. Blots were developed with Pierce ECL Western Blotting substrate (Thermo Scientific; 32209) and visualized using the GeneGnome chemiluminescence imaging system. The following primary antibodies were used for immunoblotting analysis: anti-cathepsin C (D-6) (Santa Cruz; sc-74590), anti-LYAG (Sigma-Aldrich, USA; HPA029126), anti-IBP7 (Abcam; ab171085), anti-TPP1 clone 2E12 (Millipore Sigma, US; MABN1806), anti-CO8A1 (Millipore Sigma; HPA053107), anti-CA2D1 (Thermo Fisher; MA3-921), anti-SDPR (Proteintech; 12339-1-AP), anti-LAMP1 (CD107a) (BioLegend; 328601), anti-TFEB (Santa Cruz Biotechnology; sc-166736), anti-p16 (Santa Cruz; 56330), anti-Na/K ATPase (Invitrogen; MA5-32184), anti-GAPDH (Sigma-Aldrich, G9545), and anti-β-actin (Cell Signaling; 4967). The following secondary antibodies were included: anti-rabbit IgG H&L (HRP) (Abcam; ab6802) and anti-mouse IgG H&L (HRP) (Thermo Scientific; G-21040).

### Flow Cytometry

NS and S-dox IMR-90 single cell suspension was prepared in FACS buffer (Miltenyi Biotech). Cells were incubated with anti-human LAMP-1 antibody (BioLegend; 328601) or isotype control on ice for 30 minutes (1:50). Cells were washed and resuspended in ice-cold PBS. APC-conjugated anti-mouse IgG (H+L) secondary antibody (Thermo Scientific; A-865) was incubated with the cells for 30 minutes on ice. Then, cells were washed and analyzed on the flow cytometer (MACSQuant, Miltenyi). Cell viability was determined by 7-AAD staining, and live cells were gated for downstream analysis. Data were analyzed using Flowlogic software (Miltenyi Biotech, Germany).

### Lysosomal enzyme activity

Alkaline phosphatase and hexosaminidase activities were measured using a colorimetric assay with 4-Nitrophenyl phosphate disodium salt hexahydrate (Sigma-Aldrich; N4645) and 4-Nitrophenyl N-acetyl-β-D-glucosaminide (Sigma-Aldrich; N9376) as substrates in an alkaline phosphatase buffer (100 mM NaCl; 100 mM Tris-Cl; 50 mM MgCl2; 1% Tween-20). Low serum (0.2 % FBS) media was collected from cells after 24 hours of culture, while an empty culture medium served as background control. The reactions were quenched by adding 1M sodium carbonate, and the absorbance at 405 nm in each sample was measured as a readout.

### LDH release

Lactate dehydrogenase (LDH) levels were assessed using the CytoTox96 Non-Radioactive Cytotoxicity Assay (Promega; G1780), which quantitatively measures LDH released. The baseline LDH in empty culture medium was used as a negative control. Absorbance was recorded at 490 nm using SpectraMax i3 Multi-Mode Microplate Reader (Molecular Devices, San Jose, CA, USA). Average absorbance values for the culture medium background were subtracted from all experimental values.

### Crystal violet staining for endpoint cell viability

Cells in 6-well culture plates (Greiner Bio-One; 657160) were washed twice with PBS, incubated in 4% PFA for 15 minutes at room temperature, and then stained with 0.25% crystal violet for 10 minutes. After washing with deionized water, images were captured (3 per well). Crystal violet was solubilized using 1% SDS, and the absorbance at 570 nm was quantified with a SpectraMax i3 Multi-Mode Microplate Reader (Molecular Devices, San Jose, CA, USA).

### Real-time cell viability assays

To measure the viability of NS and S-dox IMR-90 cells in real time, we utilized xCELLigence RTCA-MP (Agilent, USA), which measures cellular impedance of live cells (19). Cells were seeded on E plates (Agilent, USA), incubated overnight and treated with the compounds. After the drug addition, impedance measurements were recorded every 15 minutes. Cells treated with 0.2% Triton X-100 were used as a 100% dead cell positive control for cytotoxicity assays. All experiments were performed in at least five replicates for each dose. Changes in impedance were expressed as a cell index (CI) value, which derives from relative impedance changes corresponding to cellular coverage of the electrode sensors, normalized to baseline impedance values with medium only.

### Bleomycin mouse model of fibrosis

The C57BL/6 mice (The Jackson Laboratory, strain #000664) were used in experiments. All animal experiments adhered to the NIH Guide for the Care and Use of Laboratory Animals (National Academies Press, 2011). The use of animals was evaluated and approved under Institutional Animal Care and Use Committee (IACUC) protocols (SRF-01.8).

Mice were randomized to receive either a sham injection of saline or bleomycin (bleo) (MedChemExpress HY-17565, 2.5mg/kg), delivered via an oropharyngeal route. The 2.5 mg/kg bleo solution was prepared in sterile saline (BioWorld, 40120975-2). Approximately 10% of the animals had to be sacrificed during the following ten days due to excessive weight loss and signs of deterioration. This aligns with previous reports (20). This study excluded the bleo-treated mice that neither lost weight nor reached a humane endpoint.

Ten days after the injury, mice were randomized to create groups with similar levels of weight loss: bleo-treated control mice (saline injection); nintedanib (MedChem Express, HY-50904) (i.p. in saline, 0.54 mg/kg every other day, a total of six injections); ApiD (Axon Med Chem AXON 2500) (i.p. in saline, 2 mg/kg, daily i.p. injections for a total of eight days); and ApiD (i.p. in saline, 5 mg/kg, daily i.p. injections for a total of eight days). Mouse weights and behavior were continuously monitored during and after the treatment.

Twenty-six days after the bleo injection, mice were sacrificed, and tissue samples were collected.

### Quantitative real-time PCR from mice lung tissue

After sacrifice, heart perfusion with saline was performed, and the lungs were collected. Right lungs were flash-frozen in liquid nitrogen for RNA isolation and gene expression analysis. mRNA was isolated by mechanical homogenization in liquid nitrogen, followed by RNA extraction using the Qiagen RNA extraction kit (RNAeasy Plus Mini kit 74136) and the QIAshredder kit (Qiagen 79656). All samples were tested for RNA concentration and quality using Nanodrop (Thermo Fischer). cDNA was produced using the AzuraQuant™ cDNA Synthesis Kit (Azura Genomics AZ-1996). qPCR was performed using the following TaqMan primer probes (Applied Biosystems TaqMan gene expression assays): Rpl13a housekeeping gene – Mm05910660_g1; Cdkn2a – Mm00494449_m1.

### Histological analysis for lung fibrosis

The left lungs were preserved in 10% neutral-buffered formalin (Sigma Aldrich HT501128) overnight at 4°C before being transferred to freshly prepared 70% EtOH for storage prior to paraffin embedding. The formalin-fixed paraffin-embedded (FFPE) blocks were prepared at the Zyagen facility (San Diego, CA). Five micrometer sections were deparaffinized and stained using the Masson-trichrome staining kit (VitroVivo Biotech, VB-3016) according to the manufacturer’s instructions. Fibrosis intensity scores were determined by a certified pathologist who was blinded to the identity of the samples.

For Picrosirius red (PR) collagen staining, sections were deparaffinized, rehydrated, and treated with phosphomolybdic acid 0.2% aqueous freshly made solution for 5 minutes. After a 5-minute tap water rinse, sections were stained with 0.1% in saturated picric acid for 2 hours at room temperature. Sections were incubated for 3 minutes in 0.015N freshly prepared hydrochloric acid. Sections were rinsed in 70% ethanol for 45 seconds, rehydrated, and mounted on slides. Quantification was done using Image J software, setting the same threshold for all analyzed samples.

### Immunohistochemistry for quantification of senescence in the lung

For p21 IHC staining, FFPE sections were deparaffinized, and antigen retrieval was performed in Tris-HCl buffer (pH 10) by submerging the sections in heated buffer for 10 minutes. Endogenous peroxidases were inhibited with a 10 minute incubation in 3% hydrogen peroxide. Sections were then blocked with 5% GS (Gibco, 16210-064) in TBST buffer and stained with p21 antibodies (Thermo Fisher Scientific, 14-6715-81) at a 1:50 dilution overnight at 4°C. Afterward, sections were washed and stained with secondary goat anti-rabbit antibodies (ImmPRESS HRP goat anti-rabbit HRP IgG polymer kit, Vector Laboratories, MP-7451-15) and visualized using a DAB substrate (ImmPACT DAB substrate peroxidase kit, Vector Laboratories, SK-4105). Sections were counterstained with hematoxylin (Vector Laboratories, H-3401) for four minutes before washing, rehydrating, and mounting in Permount media (Fisher Chemical SP15-100). Imaging of the entire tissue sections was conducted using 20x magnification in the Olympus Slideview VS200 slide scanner. Five random fields of view from the fibrotic regions of the tissue were blindly chosen from each sample and evaluated for p21 intensity using ImageJ software.

## Results

### Characterization of the senescent cell surface proteome

To dissect the proteomic landscape of the senescent cell surface, we first established two models of senescence using IMR-90 human fetal lung fibroblasts. To this end, we used treatment with either doxorubicin or exposure to ionizing radiation to induce senescence, two of the most well-studied models (21). We observed an increase in CDKN2A (p16^INK4A^) expression that occurred on day 5 following the treatment, reaching significantly higher expression by day 8 and remaining stable until day 13 (**Figure 1A**). As the senescence phenotype is known to be stable up to and including day 14 after induction, we performed our proteomics analysis by treating IMR-90 cells with either doxorubicin (S-dox) or ionizing radiation (S-IR) and confirmed senescence by performing assays on day 10, followed by proteomic analysis on day 14 (**Figure 1B**). In line with previous reports, these protocols consistently induced permanent cell cycle arrest, accompanied by elevated expression of the CDKN2A and CDKN1A (p21^Cip1A^) cell cycle checkpoints (**Figure 1C**). Like the S-dox cells, S-IR also exhibited significantly higher expression of p16^INK4A^ and p21^Cip1A^; however, this increase was smaller in scale than that observed in the doxorubicin-treated cells (**Figure 1C**). High expression of the cell cycle arrest mediators p16^INK4A^ and p21^Cip1A^ corresponded with a loss of Edu-positive cells, suggesting a complete halt in cell proliferation (**Figures 1D, D-i)**. S-dox and S-IR fibroblasts displayed numerous γ-H2AX nuclear foci and a marked decrease in Lamin B1 expression, underscoring DNA damage and the loss of nuclear integrity (**Figures 1E, E-I, F)**. Additionally, senescent cells showed a prominent loss of High Mobility Group Box 1 (HMGB1) protein from the nucleus, a process known to occur during senescence that leads to the activation of pro-inflammatory signaling (22) (**Figure 1E**). None of these hallmarks of senescence were evident in NS cells.

**Figure 1.**
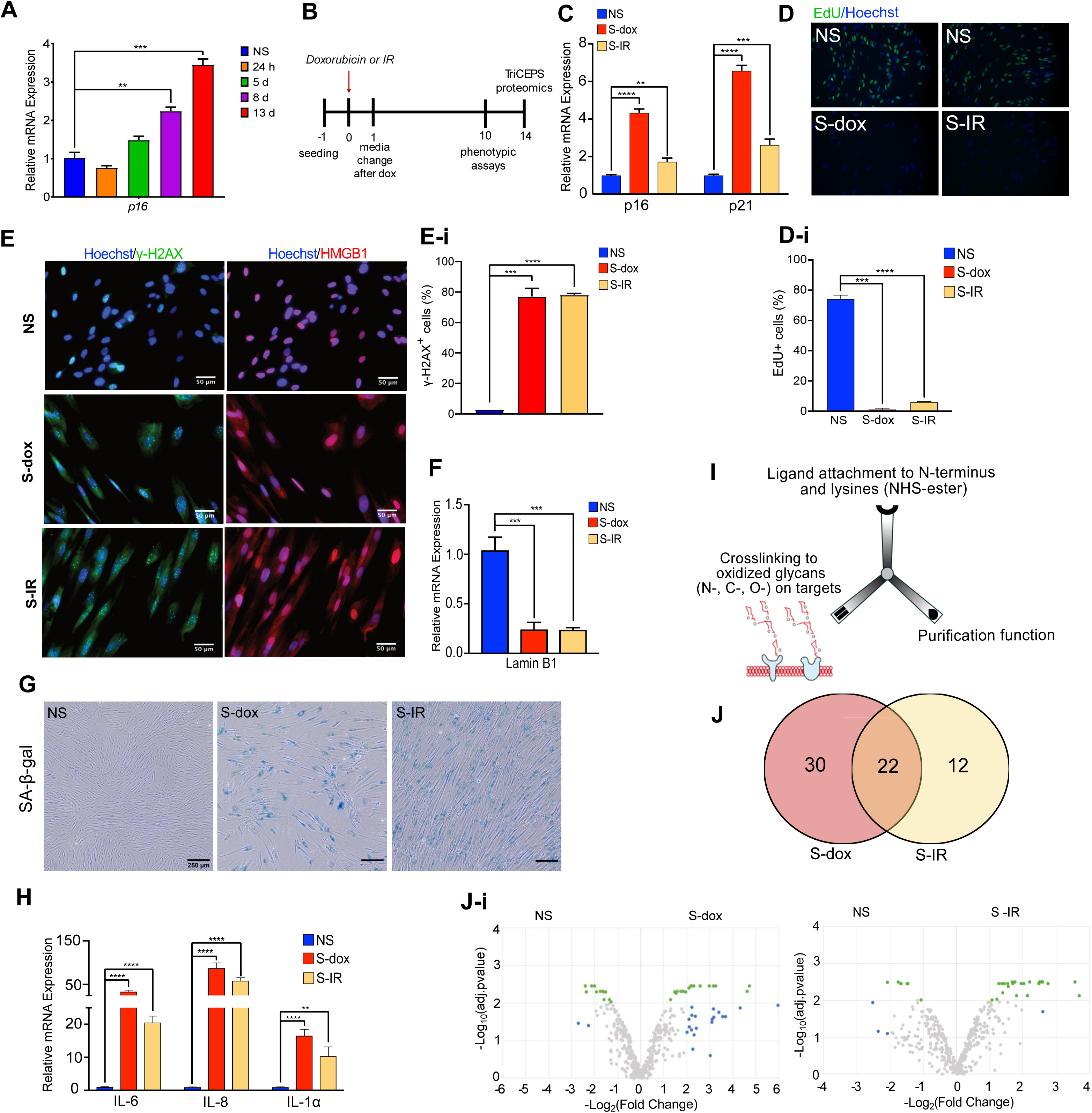
Senescent cell surfaceome profiling. **A.** Expression of CDKN2A^INK4A^ coding for p16 in IMR-90 cells at 1, 5, 8, or 13 days following treatment with doxorubicin n=3, one-way ANOVA, **p < 0.01, ***p < 0.001. **B.** Timeline of the experiments with S-dox and S-IR cells. **C.** Expression of CDKN2A^INK4A^ (p16) and CDKN1A (p21) in NS, S-dox and S-IR IMR-90 cells. One-way ANOVA, **p < 0.01, ***p < 0.001, and ****p < 0.0001; n=3. **D, D-i.** EdU-staining in NS and S-dox cells – representative image (D) and quantification (D-i). One-way ANOVA, **p < 0.01, ***p < 0.001, and ****p < 0.0001; n=3. **E, E-i.** γ-H2AX (green) and HMGB1 (red) immune-fluorescent staining in permeabilized NS, S-dox and S-IR cells. Representative images (E) and quantification of cells with two or more γH2AX nuclear foci (E-i). One-way ANOVA, ***p < 0.001, ****p < 0.0001; n=3. Scale bar = 50 μm. **F.** LaminB1 gene expression in NS, S-dox and S-IR cells. One-way ANOVA, ***p < 0.001; n=3. **G.** SA-β-gal staining in NS, S-dox and S-IR IMR-90 cells, 13 days after exposure to vehicle or doxorubicin. Representative images of >10 independent experiments, scale bar = 250 µm. **H.** Gene expression of IL-6, IL-8 and IL1-α in S-dox and S-IR IMR-90 cells. One-way ANOVA, **p < 0.01, ****p < 0.0001; n=3. **I.** Schematic of the Dualsystems Biotech AG, TriCEPS^TM^ V.3.0 compound. **J.** Venn diagram of the top proteins found as significantly enriched in S-dox and S-IR cells compared to NS cells. **J-i.** Volcano plot of Surfaceome-TriCEPS dataset comparing detected proteins in NS vs S-dox (left panel) and NS vs S-IR (right panel) cells. Proteins labeled in green as highly differentially expressed with an adjusted p ≤ 0.01. Proteins labeled in blue are highly differentially expressed with fold change greater than 4, but 0.05 ≥ p ≥ 0.01.

To further confirm senescence induction, we measured the enzymatic activity of the SA-β-gal. No staining was evident in NS cells, whereas most of the S-dox and the S-IR cells stained blue, indicating high SA-β-gal activity in acidic pH, characteristic of senescent cells in culture (**Figure 1G**). In addition, both the S-dox and the S-IR cells had significantly higher levels of expression of key components of SASP: the inflammatory cytokines interleukin-6 and-8 as well as interleukin −1 alpha (IL6, IL-8, and IL-1α, respectively) (**Figure 1H**).

Following the confirmation of robust senescence induction, we conducted a cell surface proteomic analysis utilizing Surfaceome-TriCEPS™ technology. Briefly, the hydrazone group of the TriCEPS v.3.0 reagent cross-links with oxidized, extracellularly exposed surface aldehydes of N-, C-, or O-glycosylated proteins on living cells. To perform the assay, we gently oxidized the cell surface, followed by cross-linking with the TriCEPS v.3.0 reagent. Samples were then analyzed using LC-MS/MS for unbiased detection and identification of the cell surface proteome in S-dox and S-IR cells, allowing for a quantitative comparison to NS cells (**Figure 1I**).

Using this method, we identified 52 differentially regulated surface proteins in S-dox compared to NS cells **(Supplementary table 1)**. Only 34 proteins were differentially regulated on the surface of S-IR compared to NS cells. There was a significant overlap of 22 proteins that were abundant in both the S-dox and S-IR cells, while being absent on the surface of the NS cells (**Figure 1J**). The differentially abundant proteins were categorized into high and medium-confidence groups. High-confidence differentially expressed proteins are defined as those with an adjusted p-value of less than 0.01 (-Log10(adj. p-value)>2). Medium-confidence differentially expressed proteins are those with a fold change difference greater than 4, but with statistical significance less than 0.01. These results, represented in a volcano plot **(Figure J-i),** demonstrate a change in the cell surface proteome associated with senescence.

### Senescent cells express lysosomal enzymes on the plasma membrane

To validate the results of the surfaceome proteomic screen results, we focused on the differentially regulated proteins common in both the S-dox and the S-IR (**Figure 2A**). Western blot analysis of the whole cell lysates from the NS and S-dox cells corroborated elevated amount of cathepsin C, lysosomal alpha-glucosidase, tripeptidyl peptidase 1, voltage-dependent calcium channel subunit alpha-2/delta-1 and Cavin-2, also known as Serum Deprivation Response protein (CATC, LYAG, TPP1, CA2D1 and CAVN2 respectively) proteins in the S-dox cells compared to the NS cells (**Figures 2B, B-i)**. We further confirmed the expression of these candidates in another cell type by treating human EC with doxorubicin to induce senescence, as described earlier (21). These proteins were similarly elevated in the senescent primary doxorubicin-treated EC compared to NS cells (**Figure 2C**), suggesting that cell surface expression of the identified proteins may be a common feature of senescent cells, independent of cell type of origin. In addition, these proteins were also up-regulated in the S-dox cells compared to quiescent IMR-90 that had been kept in the low serum conditions for either one (NS1) or four (NS2) days (**Figure 2D**), suggesting that a permanent cell cycle arrest and full senescence establishment is required for such protein expression, not merely a halt to cellular proliferation.

**Figure 2.**
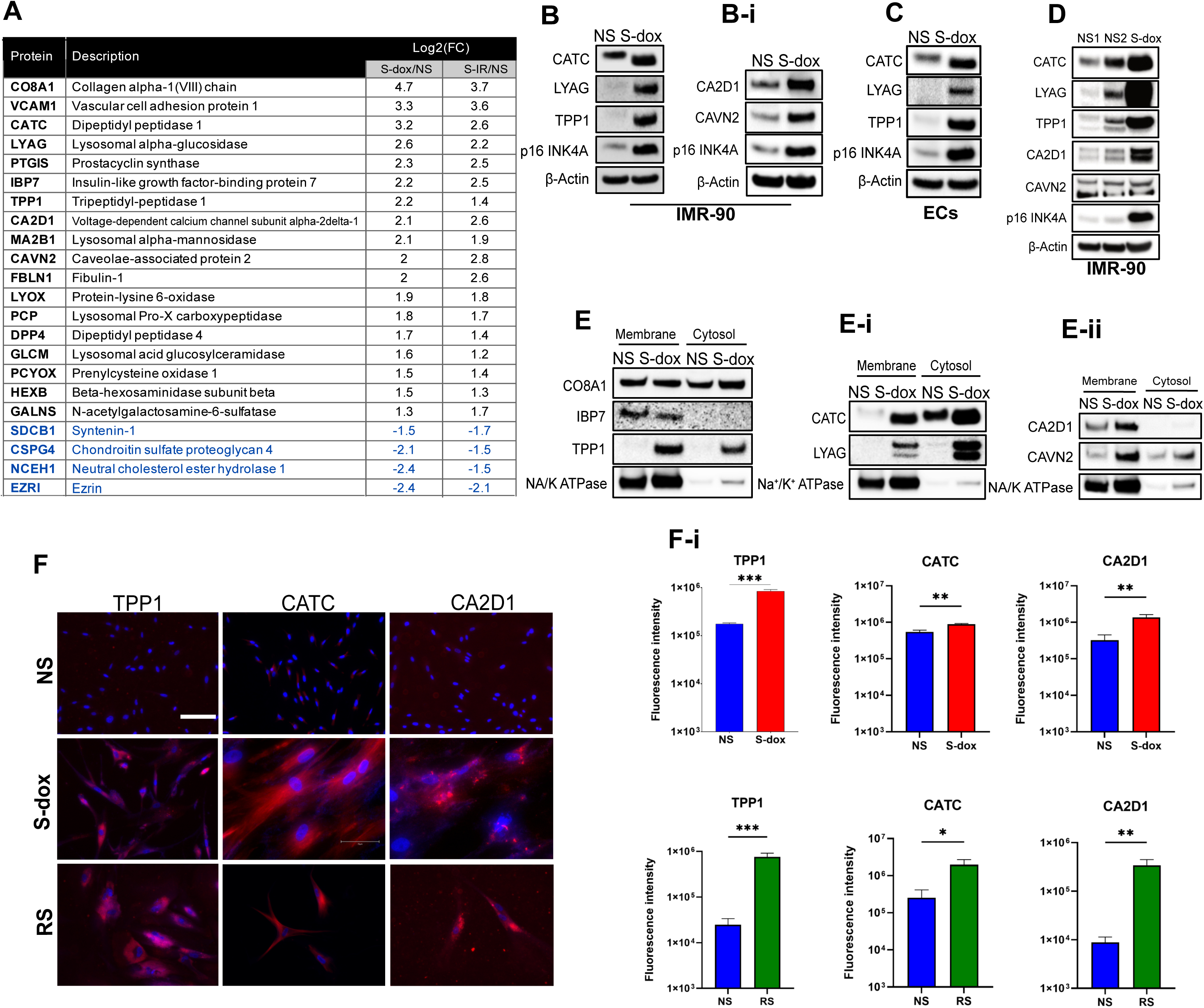
Validation of the proteomic screen results. **A.** List of the most differentially up- and down-regulated (in blue) surface proteins that were common between S-dox and S-IR cells compared to NS cells. **B, B-i.** Validation of differential protein expression of CATC, LYAG, TPP1, CA2D1 and CAVN2 in the whole cell lysate of NS and S-dox IMR-90 cells. p16 was used for validation of senescence induction, β-Actin was used as a loading control. **C.** Validation of the differential CATC, LYAG and TPP1 protein expression in the whole lysate of NS and S-dox ECs. **D.** Western blot comparing total CATC, LYAG, TPP1, CA2D1 and CAVN2 protein expression in NS cells, and IMR-90 cells kept in low serum medium for one (NS1) or four (NS2) days to induce quiescence. **E - E-i.** Western blot analysis of the CO8A1, IBP7, TPP1, CATC, LYAG, CA2D1 and CAVN2 protein isolated from the membrane and cytosolic fractions of NS and S-dox cells. Na⁺/K⁺ ATPase was used to confirm successful fraction separation. **F.** Representative immunofluorescence images of NS, S-dox and RS IMR-90 cells labelled with cell surface markers identified in the Surfaceome-TriCEPS dataset (TPP1, CATC and CA2D1) in red and Hoechst nuclear staining in blue. Scale bar = 75 µm. **F-i**. Quantification of median fluorescence intensity per cell. Multiple images were collected per sample and all clearly visible cells per sample were analyzed. Unpaired t-test, **p < 0.01, ****p < 0.0001; n≥3.

Next, we analyzed expressions of the Ctsc, Gaa, Tpp1, Cacna2d1, and Sdpr genes at various time points following the doxorubicin treatment. This revealed a time-dependent increase in the expression of Cacna2d1, while the expression of Ctsc, Gaa, and Tpp1 did not significantly change after treatment with doxorubicin. In contrast, Sdpr expression was significantly increased 24 hours after doxorubicin treatment but returned to similar levels as in the NS cells at a 5-day mark **(Figure S1A)**.

In summary, we found that CATC, LYAG, TPP1, CA2D1, and CAVN2 were elevated on the surface of senescent IMR-90 cells and primary EC, and western blot analysis revealed a corresponding increase in their levels within these cells. However, their expression was not transcriptionally upregulated. Therefore, we reasoned that selective localization enriched these proteins on the cell membrane. To confirm this, we performed a western blot analysis of the membrane and cytosolic fractions isolated from the S-dox and NS IMR-90 cells. We found that CATC, LYAG, TPP1, CA2D1, and CAVN2 were abundant in the membrane fraction collected from the S-dox cells, whereas negligible amounts were detected on the surface of NS IMR-90 cells. Notably, while CA2D1 was exclusively found in the membrane fraction, other proteins were upregulated in both the cytosolic and membrane fractions (**Figures 2E, E-i, E-ii)**. In contrast, one of the two alpha chains of type VIII collagen (CO8A1) and insulin-like growth factor-binding protein 7 (IBP7), which were among the shortlist of enriched proteins identified in the screen, did not show a significant increase in both S-dox and NS cells and were excluded from subsequent analyses (**Figure 2E**).

Upon further investigation, we examined the kinetics of protein abundance for the identified proteins on the cell surface following senescence induction with doxorubicin. We observed that the increase occurred on day five, peaking on day 13 post-treatment, which aligns with the timeline of p16 gene expression. This suggests that the expression of senescent cell surface proteins develops in parallel with the establishment of senescence and cell cycle arrest **(Figures S1B, C)**.

Finally, we validated our findings in a different mode of senescence induction: the IMR-90 cells that had approached PD numbers close to 60. We confirmed that these IMR-90 cells were in replicative senescence (RS) by measuring SA-β-gal activity. As expected, RS cells contained approximately 98% SA-β-gal-positive cells, compared to nearly none in the NS culture **(Figure S1D)**. Supporting previous results, we observed a significant upregulation of TPP1, CATC, and CA2D1 in non-permeabilized fixed S-dox and RS cells, measured using immunofluorescence (**Figures 2F, F-i)**.

The pathway enrichment analysis of the detected cell membrane proteins identifies lysosome organization and lysosome-related processes as the two most significantly upregulated pathways. This indicates that many of the identified cell surface proteins in senescent cells are enzymes that typically reside in lysosomes and perform proteolytic functions (**Figures 3A, B)**. Our earlier findings further support this, demonstrating that senescent cells upregulate the expression of lysosomal-associated membrane protein 1 (LAMP1) (12), a key regulator of lysosomal structure and function. Through immunofluorescence in permeabilized S-dox and RS cells (**Figure 3C**) and flow cytometry, we confirmed that LAMP1^+^ cells comprised approximately 75% of the S-dox cells and only about 17% of the NS cells from the total cell population (**Figure 3C-i**). Additionally, both S-dox fibroblasts and EC exhibited increased expression of transcription factor EB (TFEB), a known regulator of lysosomal biogenesis and exocytosis (**Figure 3D**).

**Figure 3.**
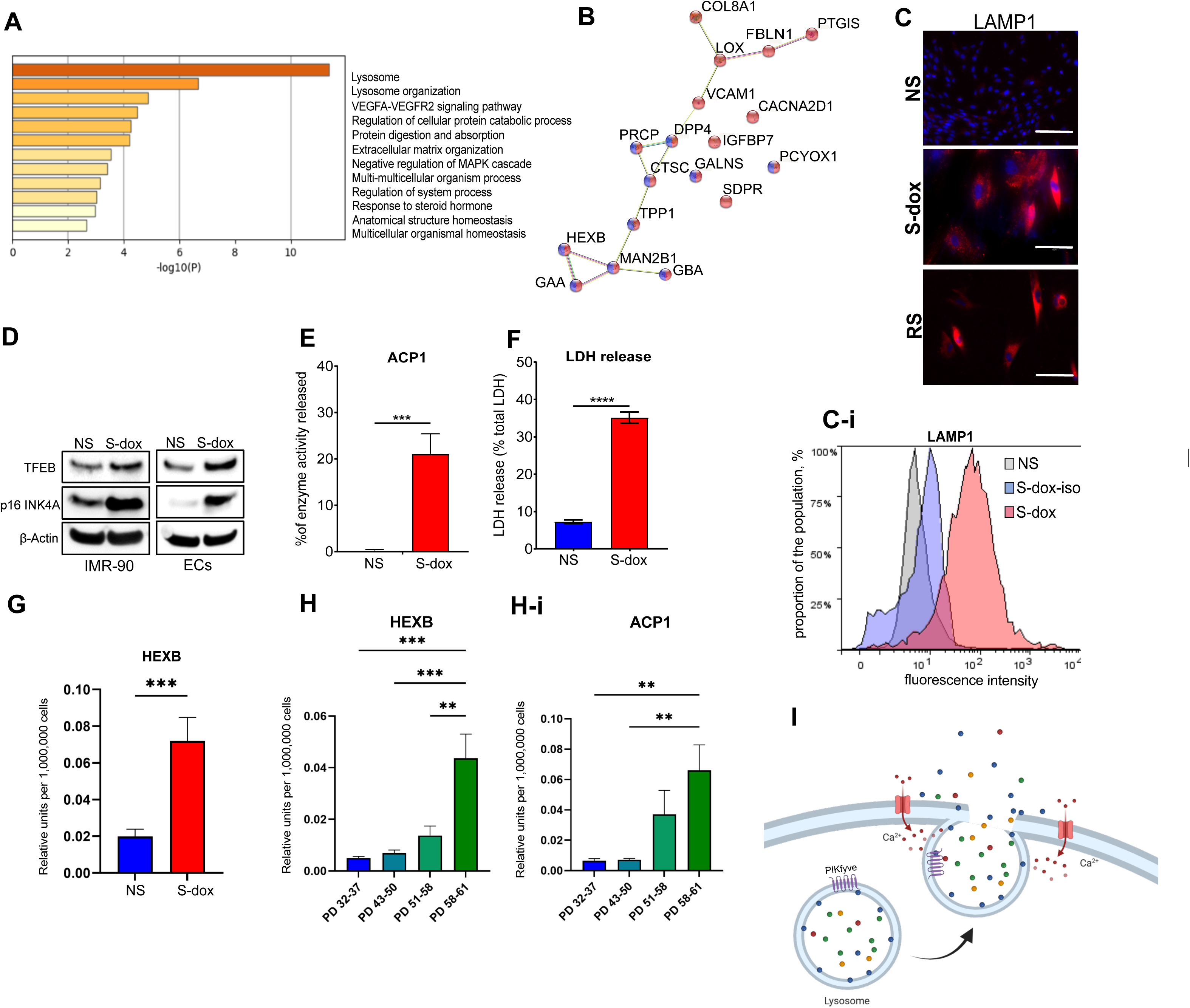
Lysosomal enzymes are exposed on senescent cell membrane via lysosomal exocytosis. **A.** Ontology enrichment clusters among the 22 proteins identified as statistically enriched, hierarchically clustered based on Kappa-statistical similarities among their gene memberships. The 0.3 kappa score was used. **B.** STRING analysis of 18 significantly upregulated senescent cell surface proteins common between S-dox and S-IR cells. Proteins identified as membrane-bound organelles in red, lysosome-associated proteins in blue. **C.** LAMP1 immunofluorescent staining in permeabilized NS, S-dox and RS cells. Representative images, scale bar = 75 µm. **C-i.** LAMP1 cell surface expression analyzed with flow-cytometry. Representative graph from ≥10 experiments. **D.** Representative immunoblots of TFEB expression in NS and S-dox IMR-90 (left panel) primary ECs (right panel). **E.** Activity of lysosomal enzyme acid phosphatase normalized to total activity in NS and S-dox cells. Unpaired t-test, ***p < 0.001; n=3. **F.** LDH release in NS and S-dox cells expressed as percentage of LDH release compared to total LDH. Unpaired t-test, ****p < 0.0001; n=3. **G.** Activity of hexosaminidase B normalized to total activity in NS and S-dox cells. Unpaired t test, ***p < 0.001; n=3. **H, H-i**. Activity of hexosaminidase B (H) and acid phosphatase 1 (H-i) normalized to the total activity in IMR-90 cells of increasing PDL values. PD 58-61 represents RS cells. One way ANOVA, **p < 0.01, ***p < 0.001; n=3. **I.** Schematic representing lysosomal exocytosis – a Ca^2+^ - mediated process of lysosomal fusion with the cell membrane that facilitates release of the lysosomal enzymes into the extracellular space.

To investigate the mechanism leading to the increase in several lysosomal enzymes and proteins on the surface of senescent cells, we measured the activity of the enzymes that are known to be released from the lysosomes into the extracellular space following cell membrane damage: acid phosphatase 1 (ACP1) and lactate dehydrogenase (LDH). We observed that both enzymes were significantly up-regulated in the conditioned media (CM) collected from the S-dox cells compared to the CM collected from NS cells (**Figures 3E, F)**. Furthermore, the enzymatic activity of the lysosomal enzyme hexosaminidase B (HEXB) in the CM from S-dox was also significantly higher than in the CM from NS cells (**Figures 3G, H, H-i)**. We also observed a progressive increase in both HEXB and ACP1 in the CM of RS IMR-90 (**Figures 3H, H-i)**. Taken together, these data suggest that senescent cells are in the state of chronic lysosomal exocytosis, which mobilizes lysosomal proteins to the cell membrane and releases lysosomal enzymes into the medium.

### PIKFyve kinase inhibition selectively targets senescent cells

Recent studies provide strong evidence that lysosomal exocytosis is central to cell membrane repair, particularly in response to mechanical or chemical damage (23–27). Because our results pointed to an elevated constitutive activity of lysosomal exocytosis in senescent cells, we speculated that senescent cells may rely on this activity to maintain cell integrity — either to repair self-inflicted cell membrane damage (suggested by the elevated level of LDH) and/or to export the toxic products of their high lysosomal number and activity. If this hypothesis were correct, then the pharmacological interruption of this activity would lead to selective senescent cell death, which could be exploited for therapeutic purposes.

To investigate if senescent cells may depend on lysosomal exocytosis for survival, we tested several small molecule inhibitors of phosphoinositide kinase, FYVE-type Zinc Finger Containing (PIKfyve kinase). PIKfyve kinase generates phosphatidylinositol-3,5-bisphosphate (PI(3,5)P₂) from phosphatidylinositol-3-phosphate (PI3P) and is a key regulator of endosomal and lysosomal membrane dynamics, including lysosomal trafficking and exocytosis, thus influencing membrane repair. PtdIns(3,5)P_2_ is required for accurate execution of the autophagic flux, delivery of the cargo to and out of the lysosomes, lysosomal fusion with the cell membrane, and the control of intracellular calcium homeostasis (28) (**Figure 3I**). PIKfyve kinase inhibitors have been reported to be far more lethal to autophagy-reliant melanoma A375 cells than the classical lysosomal inhibitors chloroquine and hydroxychloroquine (29), and the PIKfyve kinase inhibitor apilimod blocks lysosomal exocytosis and associated migratory and invasive potential in pancreatic ductal adenocarcinoma cells that rely on this pathway (30).

Our results show that treatment of S-dox cells with apilimod resulted in rapid accumulation of enlarged vacuoles (**Figure 4A**). Similar vacuoles initially appeared in the NS cells. Unlike those in S-dox, however, the vacuoles in NS cells eventually resolved. Apilimod treatment led to a dose-dependent selective killing of S-dox IMR-90 cells, with minimal effect on the viability of NS cells (**Figures 4B, B-I, C, S2B)**. We also observed that treatment with a different PIKfyve kinase inhibitor, vacuolin-1, produced similar senolytic effects on S-dox IMR-90 cells (**Figures 4D, S2A).**

**Figure 4.**
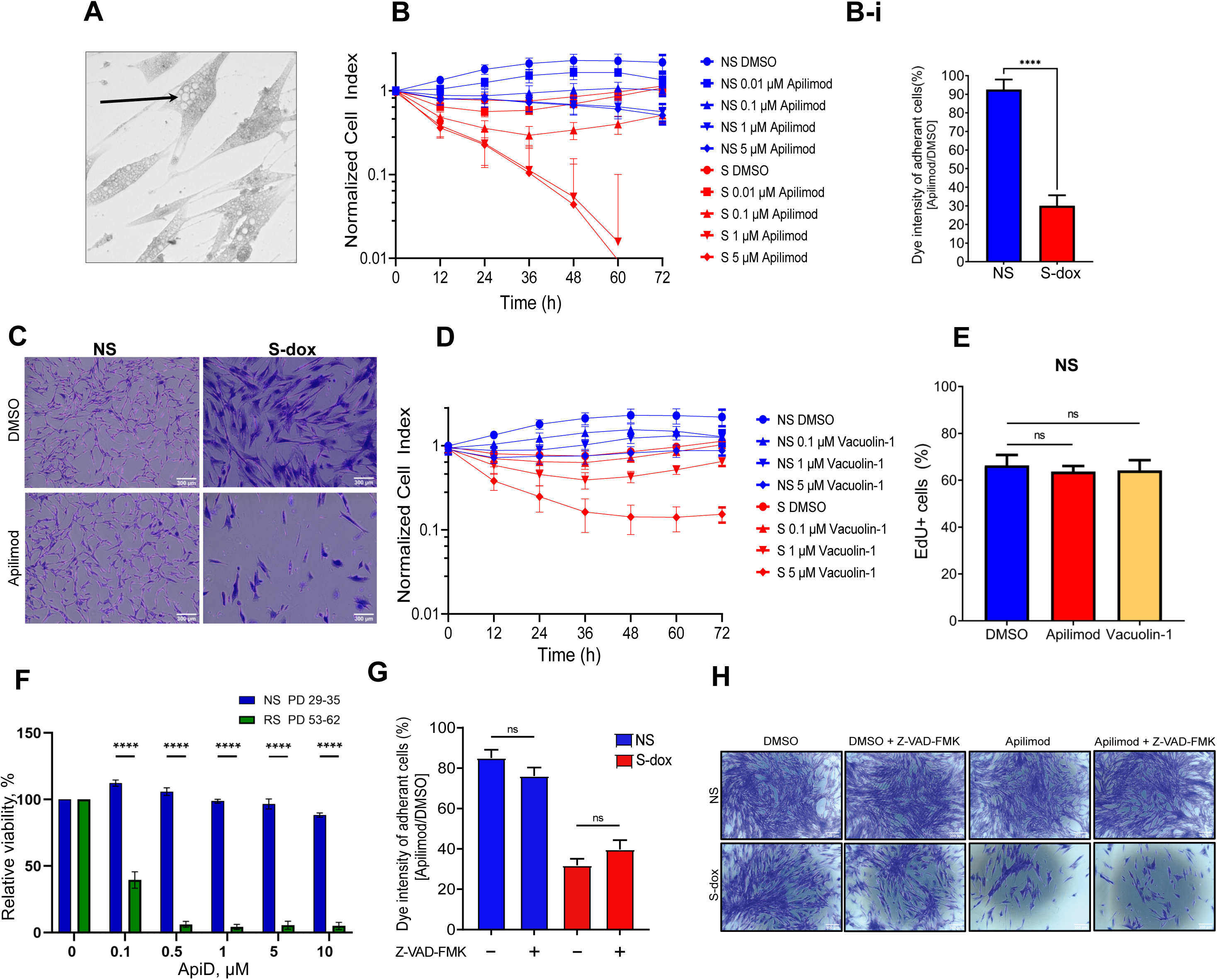
PIKfyve kinase inhibition is selectively toxic to senescent cells. **A.** A representative image of IMR-90 cells treated with apilimod 1 µM for 48 hours showing vacuolization (arrow). Cells were stained with crystal violet prior to imaging. **B.** Viability in NS and S-dox cells treated with increasing doses of apilimod over time. Impedance assay, representative results; n=3 **B-i.** Quantification of relative viability in NS and S-dox cells after 48 hours of treatment with 1 μM apilimod. Crystal violet. Unpaired t-test, ****p < 0.0001; n=3 **C.** Representative images of NS and S-dox cells treated with apilimod at 1 µM for 48 h, stained with crystal violet. Scale bar = 300 µm. **D.** Viability of the IMR-90 cells treated with increasing doses of vacuolin-1 over 72 hours measured with impedance assay. Representative experiment; n=3. **E.** Quantification of EdU incorporation in NS IMR-90 after 48 h of treatment with 1 µM apilimod. n=3. **F.** Relative viability in NS (PD 29-35) and RS (PD 53-62) IMR-90 cells after treatment with increasing doses of ApiD; n=3, unpaired t-test, ****p ≤ 0.0001. **G.** Viability of the NS and RS IMR-90 cells treated with apilimod (1 µM, 48 h) with or without pre-treatment with Z-VAD-FMK. Crystal violet assay; n=3**. H.** Representative images of the crystal-violet stained cells after treatment with apilimod (1 µM, 48 h) with or without pre-treatment with Z-VAD-FMK.

Even more striking cytotoxicity was observed when RS cells were treated with low concentrations of apilimod dimesylate (ApiD) (described below). In RS IMR-90 cells, ApiD induced nearly a complete loss of viability at a concentration of 500 nM (**Figure 4F**). In contrast, treatment with either apilimod or vacuolin-1 did not impact the proliferative potential of NS cells (**Figures 4E, S2C)**. Interestingly, apilimod did reduce the viability of quiescent IMR-90 cells maintained in low serum media for 5 to 7 days. However, this reduction was significantly milder than the cytotoxicity observed in S-dox and RS cells **(Figures S2D, D-i)**. In summary, apilimod selectively kills S-dox and RS cells while leaving the viability or proliferation of NS cells unaffected.

Next, we aimed to determine how PIKfyve inhibition influences cell death in senescent cells. Senescent cells are known to resist apoptosis (31) and the significant accumulation of enlarged vacuoles suggests that apilimod-treated senescent cells may be undergoing methuosis – a type of cell death caused by an excess of fluid-filled vesicles. To eliminate the possibility of apoptosis, we treated the cells with Z-VAD-FMK, a compound that irreversibly binds to and inactivates caspases, thus preventing apoptosis. Our findings indicate that this treatment did not prevent the loss of S-dox viability after treatment with ApiD (described below), supporting the notion that these cells are experiencing a non-apoptotic form of cell death (**Figure 4G, H)**.

### Apilimod reduces pulmonary fibrosis and decreases senescence in bleo-treated mice

After confirming that apilimod selectively eliminates senescent cells *in vitro*, we tested this intervention in a vertebrate animal model. For this purpose, we utilized ApiD, which exhibits superior pharmacokinetics due to its enhanced solubility in aqueous solutions (NCT02594384).

We initially confirmed the senolytic effect of ApiD in cell culture, noting its similarity to that of apilimod. ApiD induced abnormal vacuolization prior to cell death, akin to the parent drug **(Figure S3A)**. We also tested ApiD *in vivo* and found it to be well tolerated; three intraperitoneal injections (i.p.) of either 10, 20, or 40 mg/kg ApiD in mice exhibited no significant adverse effects **(Figure S3B)**.

We tested the efficacy of the senolytic effect of ApiD in the bleo-induced model of lung fibrosis. This treatment induces damage consistent with some features of human IPF. This age-associated interstitial lung disease is known to be associated with a high senescence burden in the lungs, and treatment with senolytic drugs has been reported to ameliorate or reverse animal models of the disease (17).

To induce IPF in mice, we treated the animals with 2.5 mg/kg bleo, delivered through an oropharyngeal route to achieve an equal and broad distribution of bleo in both lungs. Ten days after instillation the majority of mice experienced a significant reduction in body weight **(Figure S3C)**. At that point, a subset of bleo-treated mice received either ApiD (2 or 5 mg/kg, eight daily i.p. injections) or nintedanib (0.54 mg/kg, once every other day i.p. injections, total of 6) – a multi tyrosine-kinase receptor inhibitor that is standard of care for IPF. This time point has been previously reported as the point of fibrosis establishment in the lungs, which strongly correlates with weight loss (32–35). The lung tissue was harvested on day 26 to characterize fibrosis and senescence burden (**Figure 5A**). Gene expression analysis of the lung tissue revealed a significant increase in p16 expression in the bleo-treated mice compared to the sham controls. Interestingly, apilimod-treated animals had a lower level of p16 expression than the bleo-treated group, although this result did not reach significance due to the inter-mouse variation (**Figure 5B**). However, immunohistochemistry (IHC) analysis of the lung sections with p21 antibody showed a notable accumulation of p21-positive cells in the lungs of the bleo-treated mice, while there was no significant difference between ApiD-treated and sham animals (**Figures 5C, C-i)**. These data suggest that ApiD acts as a senolytic *in vivo*, clearing senescent cells from the tissue following senescence-inducing lung injury, making it a promising candidate for treatment of IPF.

**Figure 5.**
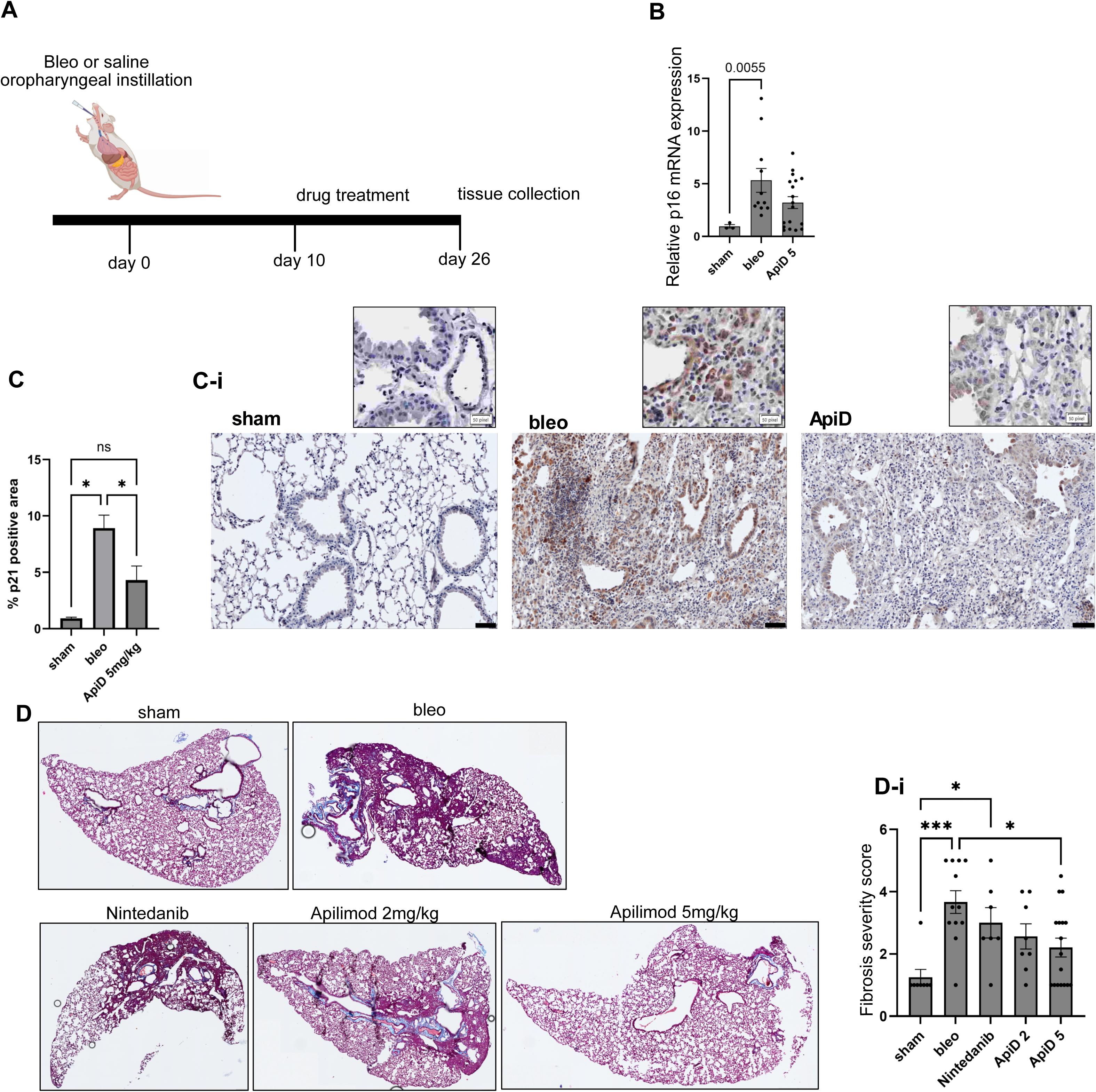
Apilimod decreases pulmonary fibrosis and reduces senescent cell burden in tissue. **A.** Schematic timeline of IPF mouse experiments. Drug treatment was initiated on day ten after bleo instillation. Tissue collection was done on day 26. Created with Biorender. **B.** Relative CDKN2A (p16) gene expression in the left lung tissue normalized to the expression in sham group. One-way ANOVA. Mice that did not lose any weight (≥100%) after bleo instillation were excluded from analysis. **C, C-i.** IHC staining and quantification of the p21 signal intensity in sham-treated, bleo-treated control, and ApiD-treated mouse lungs. Quantification of all included samples (C), One-way ANOVA, *p < 0.05. Representative images of the stained tissues (C-i). Scale bar = 50 μm. **D.** Representative images of the Masson-trichrome-stained mouse lungs collected from sham and bleo-treated animals that received either saline (bleo), or nintedanib, or ApiD (either 2 or 5 mg/kg). **D-i.** Fibrosis intensity scoring in all included samples. One-way ANOVA, *p < 0.05, ***p < 0.001; n≥7.

Furthermore, we observed a significant reduction in the severity of fibrosis in the ApiD-treated animals, as determined by a pathologist who scored the Masson-trichrome stained lung samples (**Figures 5D, D-i)**. This effect appeared to be dose-dependent, with the higher dose of 5 mg/kg producing a greater effect than the 2 mg/kg treatment. By contrast, nintedanib treatment did not significantly reduce the fibrosis score (**Figures 5D, D-i)**. In agreement with these findings, picrosirius staining of the samples also showed a significant increase in collagen deposition in bleo-treated mice. At the same time, ApiD (5mg/kg) treated animals had no significantly different level of collagen deposition compared to sham mice **(Figures S3D, D-i)**.

## Discussion

Leonard Hayflick and Paul Moorhead discovered cellular senescence in 1961, demonstrating that human fetal fibroblasts in culture possess a finite proliferative capacity (36). In recent decades, senescent cells have emerged as an attractive therapeutic target for various age-related diseases. In preclinical models, senolytic interventions have shown significant promise (37), and several senolytics have progressed into clinical development ((38, 39), NCT04129944; reviewed in (40)). One major obstacle to clinical application is the absence of a reliable and tractable biomarker for senescence. While proteomic analysis of senescent cell surfaces has been attempted previously (41), it has not yet yielded a dependable biomarker. Antibody-dependent cell-mediated cytotoxicity (ADCC) assays have previously indicated that cell surface dipeptidyl peptidase 4 (DPP4) preferentially sensitizes senescent fibroblasts, rather than proliferating ones, to cytotoxicity by natural killer cells (42), and we have demonstrated that it can be used to segregate mixed populations of senescent and non senescent cells (21). Similarly, the urokinase-type plasminogen activator receptor (uPAR) has been reported to be induced by the transition into senescence and to serve as a potential target for engineered T cell immune surveillance (43, 44). However, both of these cell-surface markers exhibit nontrivial expression in some non senescent cell types, diminishing their scientific and clinical utility.

Here we characterized the ‘surfaceome’ of senescent cells by employing a method that tags oxidized extracellular aldehydes found on the surfaces of N-, C-, or O-glycosylated proteins in living cells. This involves cross-linking them with the hydrazone group of the TriCEPS reagent, which enables subsequent affinity enrichment and MS-based analysis. This approach provides a more selective way to capture cell surface proteins (41, 42, 45).

Through this analysis we identified several previously reported senescent cell surface proteins, including CO8A1 (46), vascular cell adhesion protein 1 (VCAM1) (47, 48), and insulin-like growth factor-binding protein 7 (IBP7). IBP7 is highly expressed in senescent cells, particularly in fibroblasts, endothelial cells, and epithelial cells, and it is prevalent in the senescence-associated secretory phenotype (SASP), which contributes to chronic inflammation and fibrosis (49, 50).

We focused on the previously unreported proteins CATC, LYAG, and TPP1, which were confirmed to be elevated in multiple models of senescence induction and across various cell types. The kinetic analysis of the western blot using cytosolic and membrane fractions, along with the quantitative real-time PCR conducted during the progression of senescence induction after doxorubicin treatment, indicates that these proteins re-localize to the cell surface rather than being transcriptionally up-regulated. Metascape analysis of the proteomics data detected an enrichment of lysosomal proteins on the surface of senescent cells.

Rovira et al.’s proteomic analyses of senescent cells revealed both quantitative and qualitative changes in lysosomal resident proteins, cargo proteins delivered to lysosomes for degradation, and the secretion of lysosomal cathepsin in the SASP (51). Consistent with this report, we observed the secretion of various lysosomal enzymes.

The earliest characterization of senescent cells is based on SA-β-gal activity (52). Senescent cells are known to increase lysosomal content (53). However, their lysosomes are dysfunctional; they have a higher pH, a damaged membrane, and reduced proteolytic capacity (54). Curnock *et al.* further demonstrate that increased nuclear TFEB is necessary for senescent cell survival (54). This aligns with our observation of elevated TFEB expression in senescent cells. Interestingly, treatment with urolithin A, an inducer of TFEB activity, decreases secretion of SASP and DAMP factors by senescent cells (55).

Our earlier findings that demonstrated the accumulation of LAMP1 on the surface of senescent cells, both *in vitro* and *in vivo*, also support the increased abundance of LAMP1 in these cells. This suggests that LAMP1 may serve as a novel senescence biomarker (12). We observed elevated LDH levels in the conditioned media of senescent cells, consistent with the published literature, which may reflect a shift in cell metabolism towards aerobic glycolysis and increased lactate production in senescent stromal cells (56), but may also result from plasma membrane damage, for which lysosomal exocytosis acts as a repair mechanism (23–27).

LDH release from cells is often viewed as an indicator of membrane damage and cytotoxicity (57), while lysosomal exocytosis plays a crucial role in plasma membrane repair across all cell types (23–27). Therefore, an increase in LDH, the presence of lysosomal LAMP1 on the surface, and the secretion of lysosomal contents, including enzymes, suggest enhanced lysosomal exocytosis in senescent cells. This intracellular Ca²⁺-regulated process helps restore plasma membrane integrity after injury and contributes to regulating immune responses, bone remodeling, and extracellular matrix degradation. During lysosomal exocytosis, lysosomes located near the site of injury rapidly migrate to and fuse with the plasma membrane, effectively sealing the damaged area while releasing lysosomal contents into the extracellular space. Our findings demonstrate that lysosomal exocytosis is essential for the survival of senescent cells. Inhibition of PIKfyve kinase, a key component of lysosomal exocytosis, using vacuolin-1, apilimod, or ApiD, selectively and significantly eliminates senescent cells through non-apoptotic cell death, with minimal impact on the viability of NS or quiescent cells.

Apilimod, a potent and selective inhibitor of PIKfyve, is toxic to various cancer cells (58). Results from Phase I clinical trials (NCT02594384) indicate that apilimod is well-tolerated and provides potential therapeutic benefits for patients with non-Hodgkin’s lymphoma.

To evaluate the senolytic potential of PIKfyve kinase inhibition *in vivo*, we used a mouse model of IPF. This terminal disease, prevalent among the elderly, is characterized by an increased burden of cellular senescence, and senolytic interventions have demonstrated promise in preclinical IPF models (59). Our data confirms the senolytic activity of ApiD *in vivo*. Treatment with ApiD significantly reduced lung fibrosis in bleo-treated mice, surpassing the results observed in mice treated with nintedanib, the non-senolytic standard of care for IPF. Consistent with these findings, ApiD has previously been shown to decrease cardiac fibrosis by inhibiting TGF-β signaling (60). These results strongly support ApiD as a potential disease-modifying drug candidate for the treatment of IPF.

We recently demonstrated that senescent cells resist ferroptosis, a form of cell death, by upregulating GPX4 to reduce oxidative stress (21). Since senescent cells are also known to secrete oxidized lipids (61), it is plausible that lysosomal exocytosis plays a role in the survival mechanism by secreting oxidized lipids to diminish oxidative stress. If this is the case, then inhibiting lysosomal exocytosis could selectively increase stress on senescent cells, potentially contributing to cell death.

Our cell culture data indicate that pretreatment with the synthetic peptide z-VAD-FMK, an inhibitor of caspases that prevents apoptosis, does not block cell death induced by aplimod, thus confirming a non-apoptotic cell death mechanism. Previous studies have suggested that the eventual outcome of cellular senescence may be methuosis (62), a distinctive form of non-apoptotic cell death characterized by the accumulation of large cytoplasmic vacuoles (63). Our findings support the notion that apilimod-induced senolysis is accomplished via methuosis. In agreement with that, PIKfyve kinase, the target of apilimod, has been identified as a target of methuosis inducers (64–66). Furthermore, our light microscopy imaging of cells treated with aplimod showed accumulation of large cytoplasmic vacuoles, which resolve in non-senescent cells but lead to cell death in senescent ones.

In summary, the discovery of senescent cells’ critical role in aging and age-related diseases has significant implications for future medical treatments. These cells not only contribute to the aging process but also present an emerging target for therapeutics aimed at mitigating disorders in which they are implicated, such as IPF. Our findings reveal that senescent cells depend on lysosomal exocytosis for their survival, which opens promising avenues for therapeutic intervention using a PIKfyve inhibitor, such as apilimod. This connection underscores the potential of apilimod and its derivative, ApiD, as powerful senolytics, warranting further investigation. Such research could shift treatment paradigms for aging-related conditions, ultimately improving patient outcomes and quality of life.

## Acknowledgements

The authors would like to thank Esmeralda Jimenez for her help with animal maintenance, welfare checks and assistance with mouse experiments. We would also like to acknowledge Ashley Brauning for conducting training and providing technical assistance. Finally, we extend our gratitude to the donors of Lifespan Research Institute (formerly SENS Research Foundation).

## Author contributions

AB, KK, AS and GML performed experiments for the study and analyzed the data. A. Sharma and AB wrote the manuscript; MR provided critical discussion and extensively edited the manuscript; A. Sharma designed and supervised and obtained funding for the project; CJLS provided pathology analysis and critical comments; All authors reviewed and edited the final manuscript. All authors have read and agree to the published version of the manuscript.

