## Supplemental Figures for "Inhibition of PIKfyve kinase induces senescent cell death by suppressing lysosomal exocytosis and leads to improved outcomes in a mouse model of idiopathic pulmonary fibrosis"

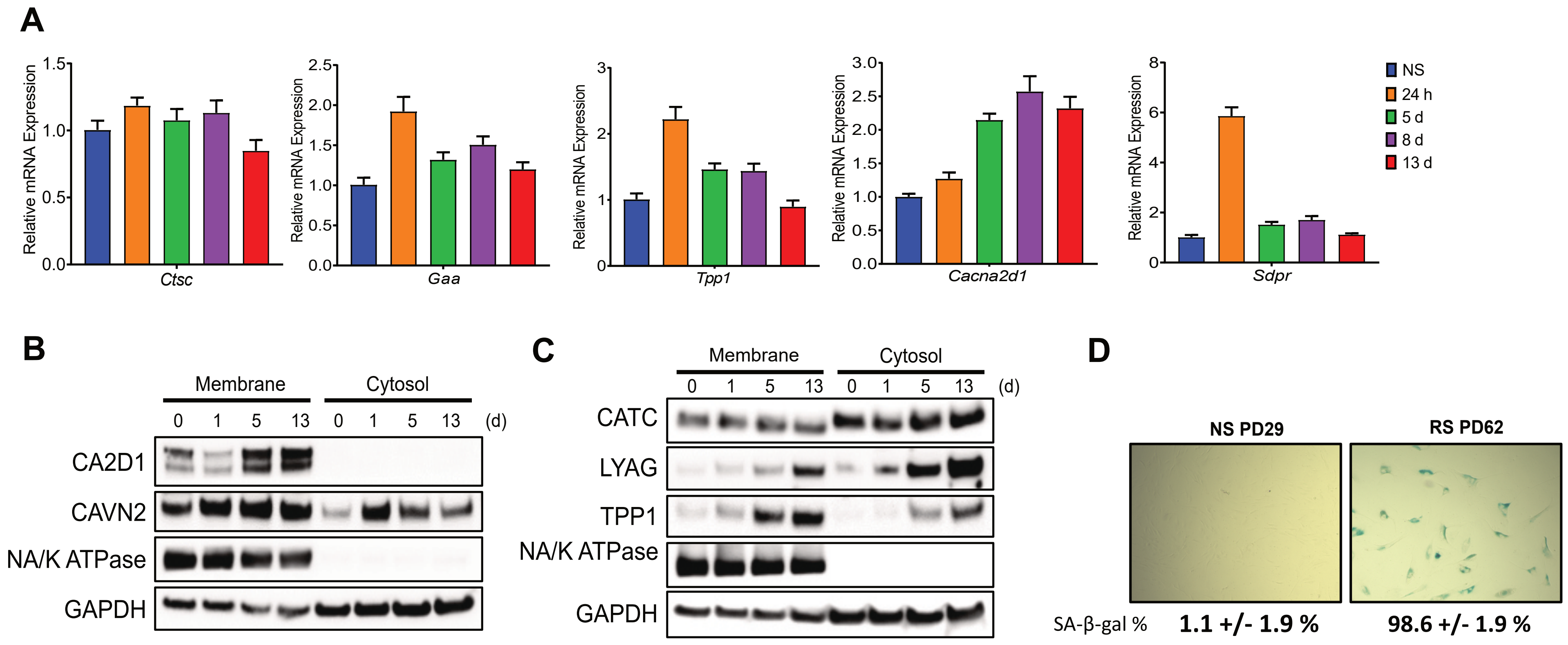
A
Relative mRNA Expression
Ctsc
Gaa
Tpp1
Cacna2d1
Sdpr
NS
24 h
5 d
8 d
13 d
B
Membrane
Cytosol
0 1 5 13 0 1 5 13 (d)
CA2D1
CAVN2
NA/K ATPase
GAPDH
C
Membrane
Cytosol
0 1 5 13 0 1 5 13 (d)
CATC
LYAG
TPP1
NA/K ATPase
GAPDH
D
NS PD29
RS PD62
SA-β-gal %
1.1 +/- 1.9 %
98.6 +/- 1.9 %

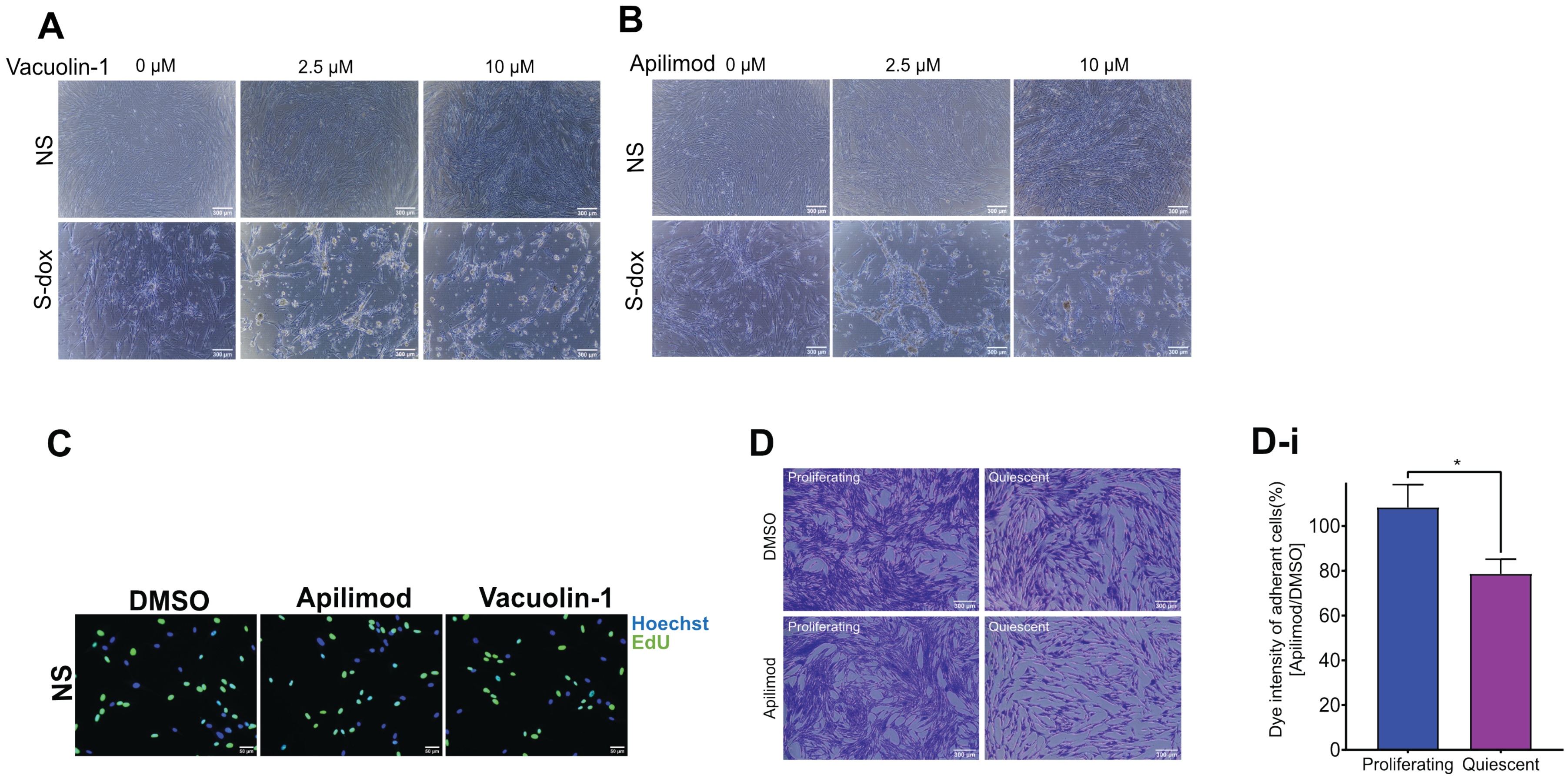
A
Vacuolin-1
0 μM
2.5 μM
10 μM
NS
S-dox
B
Apilimod
0 μM
2.5 μM
10 μM
NS
S-dox
C
DMSO
Apilimod
Vacuolin-1
Hoechst
EdU
NS
D
DMSO
Apilimod
Proliferating
Quiescent
Proliferating
Quiescent
D-i
Dye intensity of adherant cells(%)
[Apilimod/DMSO]
0
20
40
60
80
100
*
Proliferating
Quiescent

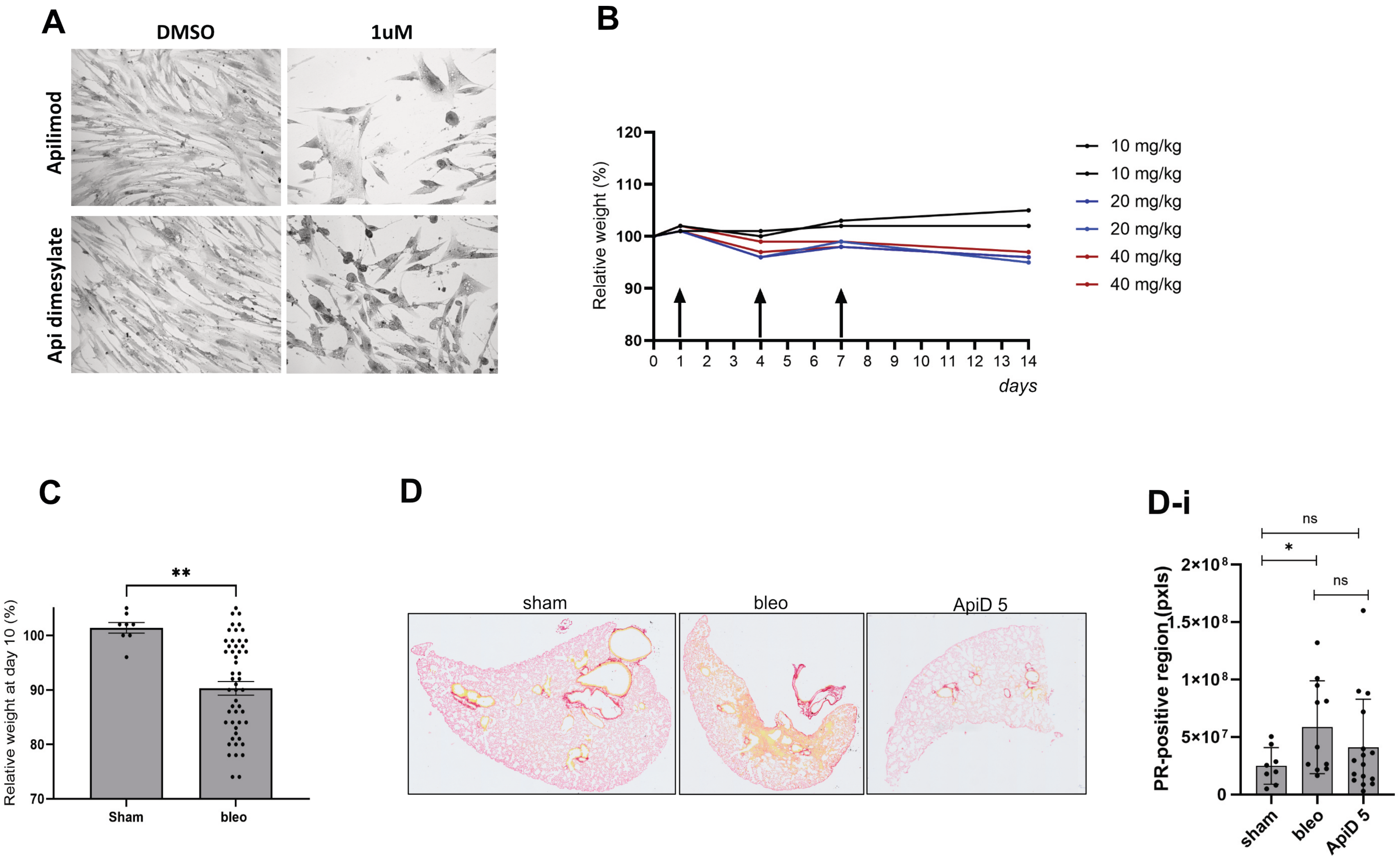

A
DMSO
1uM
Apilimod
Api dimesylate
B
Relative weight (%)
120
110
100
90
80
0 1 2 3 4 5 6 7 8 9 10 11 12 13 14
days
10 mg/kg
10 mg/kg
20 mg/kg
20 mg/kg
40 mg/kg
40 mg/kg
C
**
Relative weight at day 10 (%)
100
90
80
70
Sham
bleo
D
sham
bleo
ApiD 5
D-i
ns
*
ns
PR-positive region (pxls)
2×10⁸
1.5×10⁸
1×10⁸
5×10⁷
0
sham
bleo
ApiD 5

**Figure S1.**

**A.** Time course of relative mRNA expression of *Ctsc* (CATC), *Gaa* (LYAG), *Tpp1* (TPP1), *Cacna2d1* (CA2D1) and *Sdpr* (CAVN2) in NS and S-dox IMR-90 cells, 0 (NS) to 13 days after doxorubicin treatment. **B, C.** Time course of protein expression of top five differentially expressed proteins in the screen, from 0 to 13 days after doxorubicin treatment in membrane and cytosolic lysates of NS and S-dox IMR-90 cells. $Na^+/K^+$ ATPase and GAPDH were used to confirm cell fractionation. **D.** Representative images of NS (PD29) and RS (PD 62) IMR-90 cells stained using the SA-β-gal detection kit.

**Figure S2.**

**A, B.** Representative bright-field images of NS and S-dox IMR-90 cells after 72 hours of treatment with vacuolin-1 (A) or apilimod (B). Scale bar – 300 μm. **C.** Representative images of NS cells treated with apilimod or vacuolin-1 for 72 hours (EdU – green). Scale bar – 50 μm. **D, D-i.** Crystal violet staining (D) and quantification of proliferating NS and quiescent IMR-90 (48 hours of low serum media) cells treated with 1 μM apilimod for 48h.

**Figure S3.**

**A.** Crystal violet-stained S-dox cells treated with 1 μM of either free-base apilimod or apilimod dimesylate for 72 hours. Representative images. **B.** Relative weight change in the mice injected with 10, 20 or 40 mg/kg ApiD. 3 separate injections on days indicated with arrows. **C.** Relative mouse weight, normalized to the weight on the day of bleo injection in sham and bleo-treated mice; unpaired t-test, **$p<0.01$. **D.** Representative images of the whole lung sections stained with PR. **D-ii.** Quantification of PR staining showing PR-positive region in pixels; one-way ANOVA, *$p<0.05$.
